# Gibberellin response in the embryo epidermis regulates germination uniformity in response to seed priming

**DOI:** 10.1101/436121

**Authors:** Jack Mitchell, Nur K. Mukhtar, Isobel Skinner, George W. Bassel

**Affiliations:** School of Biosciences, University of Birmingham, Birmingham B15 2TT, UK

**Keywords:** Seed germination, seed quality, seed enhancement, seed priming, gibberellin, epidermis, seed uniformity, transgenic plant

## Abstract

Uniformity in seed germination remains a primary objective in plant-based food production systems, ensuring predictable and synchronized harvest dates, while suppressing weeds. Treatments including priming can be used to increase germination uniformity and increase the value of commercial seeds. Despite the economic and agronomic importance of seed enhancement treatments, little is known as to how they work at a mechanistic level. Using a combination of molecular genetics and microscopy, we established that hydropriming limits embryo growth genetic programs at an early stage of germination. Conversely, gibberellin (GA) and abscisic acid (ABA)-associated molecular processes progress to later stages of this developmental chronology. The response to GA was specifically affected in the epidermis of germinating embryos in response to hydropriming based on reporter gene expression. The reduction of GA response specifically in the embryo epidermis resulted in increased uniformity of seed germination following hydropriming relative to control seeds. This represents the identification of both a molecular signalling pathway and cell type that are acting to enhance the agronomic germination properties of seed populations. This provides molecular and cellular targets which may be genetically manipulated to enhance seed germination and food production in agronomic species.

## INTRODUCTION

Seeds enable plants to move through both time and space, determining where and when plants are established (Koornneef *et al.*, 2002). The seed to seedling transition is therefore one of the major shifts in the plant life cycle (Finch-Savage and Leubner-Metzger, 2006; Springthorpe and Penfield, 2015).

Central to the control of the timing of the induction of germination is the role of the antagonistically acting hormones abscisic acid (ABA) and gibberellic acid (GA), which inhibit and promote this transition, respectively (Finkelstein *et al.*, 2008; Holdsworth *et al.*, 2008). The hormone balance theory which describes this relationship provides a molecular thresholding mechanism by which the development state of seeds is defined (Karssen and Lacka, 1986). The most abundant hormone is thought to define whether or not a seed transitions to the germination program (Bradford and Trewavas, 1994). Each biosynthetic (Olszewski *et al.*, 2002; Seo and Koshiba, 2002) and signalling pathways for ABA (Park *et al.*, 2009) and GA (Lee *et al.*, 2002; Murase *et al.*, 2008) have been identified, enabling the mechanistic regulation of the molecular agents which control dormancy and germination to be investigated. Following the decision to germinate, the embryo within a seed commences growth (Koornneef *et al.*, 2002). This transition into a seedling is principally driven by cell expansion, rather than cell division (Bassel *et al.*, 2014; Sliwinska *et al.*, 2009). This discrete induction of growth following the initiation of the germination program is promoted by GA (Groot and Karssen, 1987; Koornneef and Van der Veen, 1980) and its induction of gene expression associated with cell wall remodelling proteins which facilitate cell growth (Dekkers *et al.*, 2013; Nakabayashi *et al.*, 2005; Narsai *et al.*, 2017). These cell expansion-associated genes may be considered the downstream targets of the germination process in light of the central role they play in the regulation of embryo growth (Bassel, 2016).

Spatially distinct domains of gene expression programs have been identified within the germinating *Arabidopsis* embryo using both gene expression analysis (Dekkers *et al.*, 2013) and the microscopic analysis of specific reporter constructs (Bassel *et al.*, 2014). The cellular sites of ABA and GA response and metabolism (Topham *et al.*, 2017), and growth-promoting cell wall-associated gene expression (Bassel *et al.*, 2014) have also been defined at single cell resolution. In non-germinating *Arabidopsis* embryos, the radicle was found to be enriched for both ABA and GA-associated synthesis and response components, leading to the proposal that this subdomain of the embryo acts as a decision-making centre in the control of seed dormancy (Topham *et al.*, 2017). Examination of gene expression associated with cell wall-associated gene expression revealed this to be first induced within the embryo radicle (Bassel *et al.*, 2014). These results collectively suggest that the radicle is where germination is initiated, a spatial site that overlaps with the decision-making centre.

The germination of seeds from an individual mother plant is typically non-uniform, with bet-hedging strategies being implemented (Bradford, 2002; Mitchell *et al.*, 2016; Rowse and Finch-Savage, 2003; Springthorpe and Penfield, 2015). This strategy is believed to improve plant fitness, while mechanisms underpinning this bet-hedging behaviour have been proposed (Johnston and Bassel, 2018).

In the context of food production systems, this bet-hedging trait is not favourable. Uniformity is a key objective in field-based agriculture at all stages of these systems. In order to achieve this, the germination of seeds must be synchronous once they have been planted. This co-ordinated crop establishment leads both to the suppression of weeds, and uniformity of the final product at harvest (Finch-Savage and Bassel, 2015).

In light of this important role for uniformity in seed behaviour, procedures have been developed by commercial seed vendors which increase this population trait (Paparella *et al.*, 2015). In a process termed “seed priming”, seeds are held in sub-optimal conditions for extended periods to repress germination (Finch-Savage *et al.*, 2004). Following the release of the seeds from this inhibitory treatment, the resulting germination profile of the seeds is more uniform. The mechanisms by which priming acts remain poorly understood, with protocols largely focusing on the efficacy of treatments rather than the mechanisms underpinning them. This limits the potential to enhance seed quality using these approaches.

It is thought that placing seeds under stress conditions during a priming treatment takes the population to a common stage along the germination process, where the stress-induced block acts to suspend their progression. Once released from this block, seeds in the population then complete germination more uniformly given that the implementation of the bet-hedging has been diminished.

Priming treatments may include unfavourably high temperature, osmotic stress, cold stress, application of chemicals, and others (Paparella *et al.*, 2015). While the placement of seeds into unfavourable conditions triggers ABA-mediated suppression of germination (Toh *et al.*, 2008), this does not provide insight into the stage at which priming is blocking germination, nor the mechanism by which this is acting to synchronize the behaviour of populations. Despite the central importance of priming in the seed industry and food production, the molecular and cellular mechanisms by which this acts to enhance seed germination uniformity remain unknown.

In this study we examined the spatiotemporal gene expression events underpinning the seed to seedling transition in *Arabidopsis*. We then compared these dynamic changes with those that occur in a seed that has been hydroprimed in an effort to provide insight into the mechanism by which this treatment is acting within seeds.

## RESULTS

### Establishment of a hydropriming protocol for Arabidopsis seeds

Priming protocols have been developed for a variety of horticultural crop species (Finch-Savage *et al.*, 2004; Finch-Savage and Bassel, 2015). These aim to increase the uniformity with which populations of seeds germinate as a primary goal, followed by the speed at which this occurs (Paparella *et al.*, 2015).

Hydropriming involves imbibing seeds in water under conditions which do not permit the emergence of the radicle. We developed a hydropriming protocol for *Arabidopsis*, which lacks any established method for seed enhancement. Seeds were imbibed at three different temperatures: 10 °C, 16 °C and 22 °C for four different periods of time: 6 h, 18 h, 30 h and 42 h. Following this controlled imbibition, seeds were dried and the resulting germination profiles were statistically analysed for speed and uniformity (See Materials and Methods).

The time at which seeds began to germinate (t10) was improved by all priming treatments (Fig. 1A and Supplementary Figure 1D). The reduction in the time it took for seed populations to reach 95% germination (t95) was more pronounced. The t95 was shortest across all temperature treatments that lasted for 30 h (Fig. 1B and Supplementary Figure 1E). This time then increased slightly at 42 h of hydropriming at all temperatures, indicative of a reduction in seed performance due to “over-priming”. This observation suggested that 30 h provides the optimum length of time to achieve increases in germination speed.

**Figure 1.**
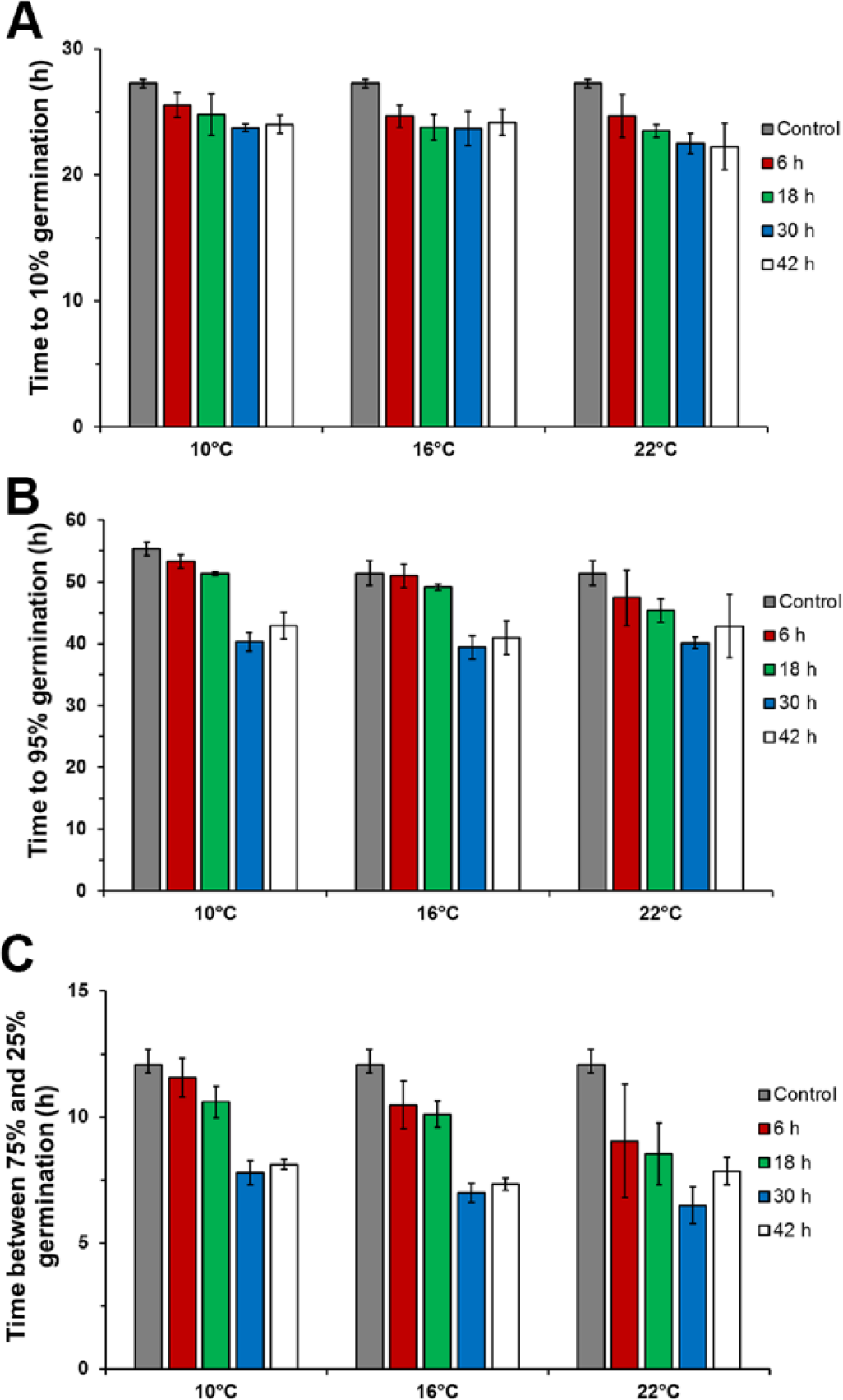
Development of an optimized priming protocol for *Arabidopsis* seeds. Seeds were primed at three temperatures: 10 °C, 16 °C and 22 °C for four different periods of time: 6 h, 18 h, 30 h and 42 h. (A) Time to reach 10 % germination (T10), (B) time to reach 95 % germination (t95) and (C) uniformity in germination for each of the treatments based on time taken between achieving 25% and 75% germination (U7525). Error bars are SD with n = 3.

The uniformity of germination in control and primed seeds was established by calculating the time taken between when the population of seeds achieved 25% to 75% germination (U7525). Again a marked improvement in uniformity was observed at 30 h of hydropriming at all temperatures examined (Fig. 1C and Supplementary Figure 1F). A slight decrease in this measure was also seen following 42 h, indicating a decrease in the beneficial impact of priming following this length of time in these conditions.

Temporally, 30 h provided the greatest enhancement in terms of germination speed and uniformity. Beyond this time, germination performance began to decrease. Similar results at this time were observed for the three temperatures observed. The hydropriming of seeds for 30 h at 22 °C was selected as a protocol to enhance the germination properties of *Arabidopsis* seeds.

These results demonstrate the ability to enhance the synchronization and speed of germination in *Arabidopsis* using hydropriming, and provide a tool to further investigate the molecular basis of seed enhancement.

### *Visualizing molecular dynamics within germinating* Arabidopsis *embryos*

Underpinning the transition from seed to seedling is a sequence of dynamic molecular events which unfold with the germinating embryo. These dynamic gene expression and epigenetic changes have been characterized previously on a genome-wide scale (Dekkers *et al.*, 2013; Nakabayashi *et al.*, 2005; Narsai *et al.*, 2017).

We sought to build upon this work by investigating the spatiotemporal dynamics of candidate genes using reporter constructs and microscopy. The visualization of key reporters across the developmental sequence from seed to seedling enables the dynamic processes underpinning this transition to be understood. Comparing the expression pattern of these reporters in seeds which have been subjected to hydropriming may therefore provide insight into the stage at which the treatment is arresting this dynamic germination program.

Three major classes of reporter were selected. The first represent genes which encode proteins targeted to the cell wall, and promote cell expansion. The other two classes represent genes and proteins associated with ABA and GA synthesis, perception and response.

The optical heterogeneity of mature *Arabidopsis* embryos makes it not possible to visualize fluorescent proteins deep within samples (Moreno, 2006). In order to achieve this, samples can be clarified and the e-glucuronidase (GUS) reporter (Jefferson, 1989) observed throughout all cells of the tissue (Truernit *et al.*, 2008). We made use of this system to examine the spatial and temporal changes which occur in germinating *Arabidopsis* seeds.

### Cell growth-associated gene expression during the seed to seedling transition

Gene expression for enzymes that promote cell wall weakening act as a proxy to understand which cells are having their growth promoted (Cosgrove, 2005). These downstream targets of the germination process have been demonstrated to play a regulatory role in the control of this transition (Lü *et al.*, 2013). Proteins encoded by *EXPANSIN* (*EXPA*) and *XYLOGLUCAN ENDOTRANSGLUCOSYLASE/HYDROLASE* (*XTH*) genes promote cell expansion in *Arabidopsis* cells (Vissenberg *et al.*, 2005b). We examined the activity of the promoters encoding members of the *EXPA* and *XTH* multigene families that are conditionally unregulated in response to the induction of the germination program (Bassel *et al.*, 2008; Dekkers *et al.*, 2013; Nakabayashi *et al.*, 2005).

The spatiotemporal patterns of gene expression associated with growth-promoting cell wall modifying gene expression may serve as a proxy to understand spatial control of cell expansion (Cosgrove, 2005). A series of expansin genes are induced in germinating *Arabidopsis* embryos (Nakabayashi *et al.*, 2005) and promoter-GUS reporters to these genes have been generated previously (Bassel *et al.*, 2014; Stamm *et al.*, 2017). These lines were examined using light microscopy over the time course of seed germination and early seedling establishment to explore the genetic control of cell expansion. *EXPA1::GUS* is first induced in cells of the embryonic radicle, and spreading progressively up the axis and into the cotyledons during germination (Figs. 2A-F). Following germination, promoter activity is increasingly focused to the root tip and cotyledons, being excluded from the majority of the root and hypocotyl (Figs. 2G-K).

**Figure 2.**
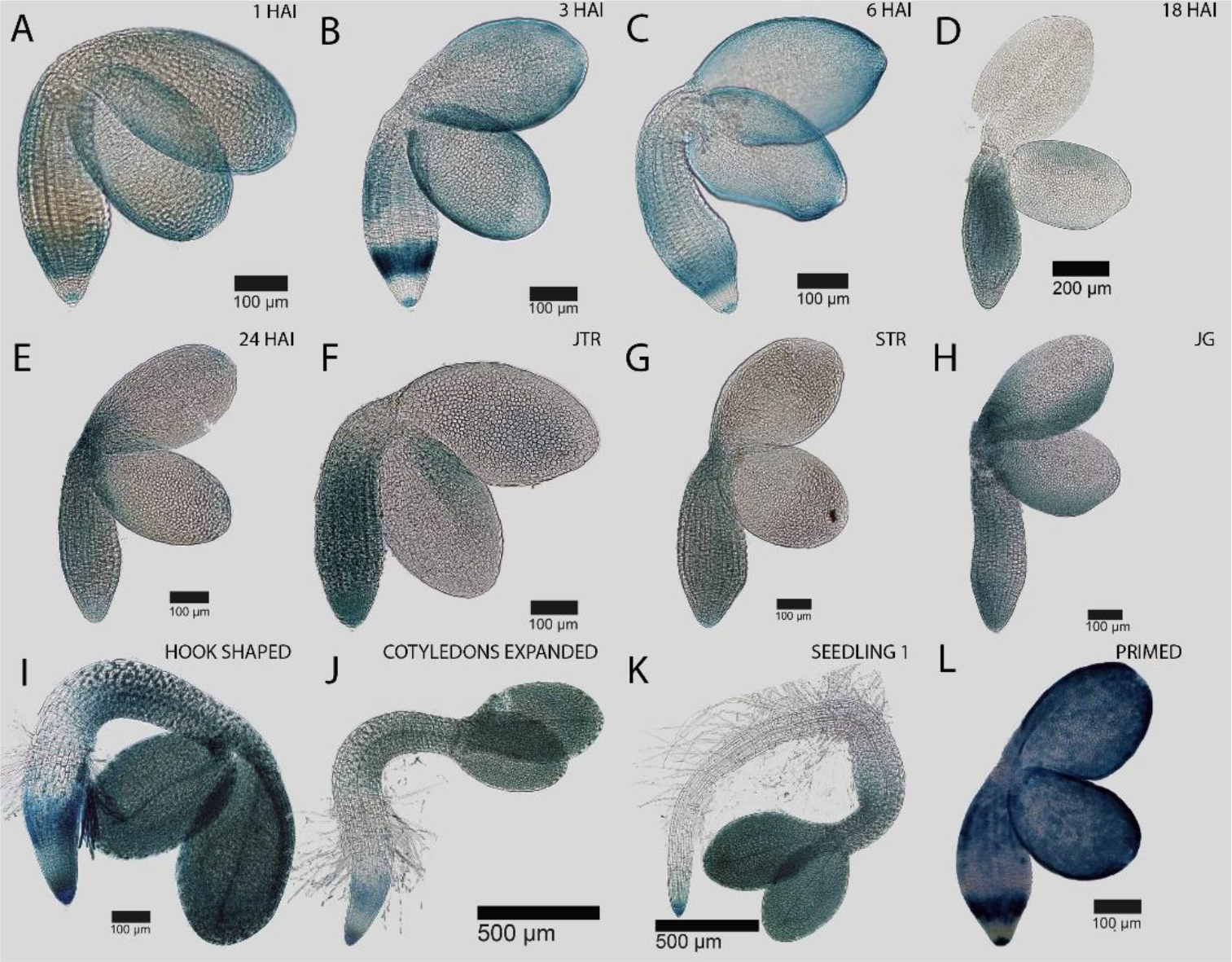
Spatial and temporal dynamics of the *EXPANSIN 1 EXPA1::GUS* promoter reporter activity during the seed to seedling transition in *Arabidopsis.* GUS activity in the germinating embryo at (A) 1 HAI, (B) 1 HAI, (C) 3 HAI, (D) 6 HAI, (E) 18 HAI, (F) early testa rupture, (G) late testa rupture, and germinated seedlings just after the completion of germination (H), (I) hook stage seedling, (J) recently expanded cotyledons and (K) a fully established seedling. (L) Pattern of *EXPA1::GUS* promoter activity in an embryo from a seed that was hydroprimed for 30 h at 22 oC. Black bars indicate the scale in each image.

Following imbibition at 22 °C for 30 h, seeds were dried and re-imbibed to examine the spatial distribution of reporters following this treatment. The expression pattern of *EXPA1::GUS* in embryos from hydroprimed seeds most closely resembled the pattern observed at 3 HAI and 6 HAI (Fig. 2L). This suggested that the block imposed by hydroprimed seeds acts at an early stage in the induction of the *EXPA1* gene. Similar results were observed for *EXPA9::GUS* (Supplementary Figure 2), *EXPA10::GUS* (Supplementary Figure 3) and *EXPA15::GUS* (Supplementary Figure 4), where primed seeds showed reporter pattern activity similar to this early stage (before 6 HAI) of the germination process.

*XTH* genes also play a role in cell wall remodelling, facilitating the expansion of plants cells (Vissenberg *et al.*, 2005a). Like *EXPA* genes, members of this gene family are upregulated during seed germination (Nakabayashi *et al.*, 2005). The activation of the *XTH19* promoter during germination initially occurs in the radicle and junction between the cotyledons and hypocotyl (Figs. 3A-C). Over the course of germination this reporter progressively moves along the hypocotyl, and into the cotyledons (Figs. 3D-H). Following germination the activity of the *XTH19* promoter becomes focused to the root and lower hypocotyl (Figs. 3H-K).

**Figure 3.**
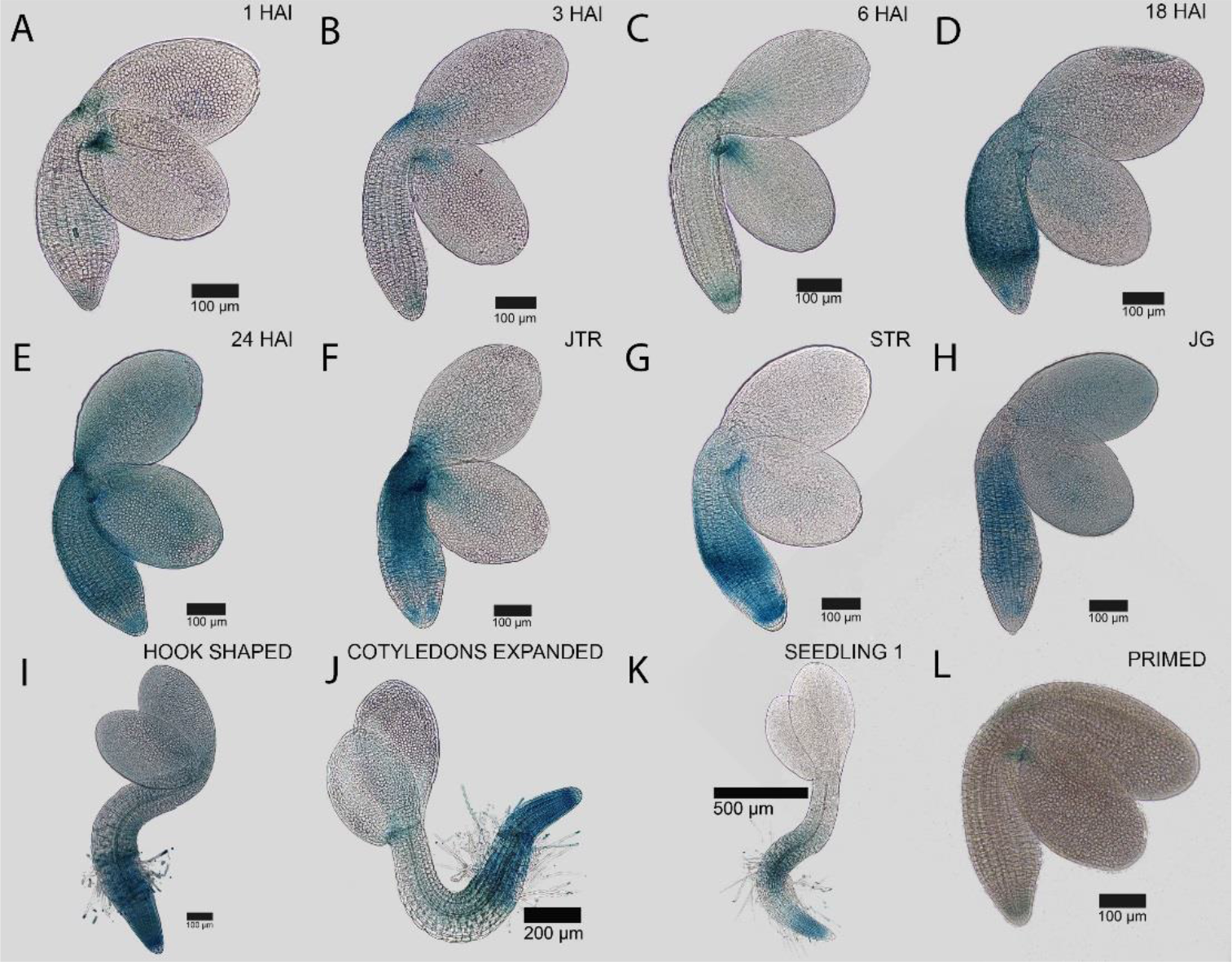
Spatial and temporal dynamics of the *ENDOTRANSGLUCOSYLASE/HYDROLASE 19*, *XTH19::GUS* promoter reporter during the seed to seedling transition in *Arabidopsis.* Promoter activity in the germinating embryo at (A) 1 HAI, (B) 1 HAI, (C) 3 HAI, (D) 6 HAI, (E) 18 HAI, (F) early testa rupture, (G) late testa rupture, and germinated seedlings just after the completion of germination (H), (I) hook stage seedling, (J) recently expanded cotyledons and (K) a fully established seedling. (L) Pattern of *XTH19::GUS* promoter activity in a hydroprimed embryo that was hydroprimed for 30 h at 22 oC. Black bars indicate the scale in each image.

Hydroprimed embryos showed an *XTH19::GUS* pattern similar to that observed at 3 HAI and 6 HAI (Fig. 3L). A similar result was observed with the *XTH18::GUS* reporter (Supplementary Figure 5), where this reporter is not induced in treated seeds, nor in the early stages of germination.

### ABA synthesis and signalling during the seed to seedling transition

ABA plays a central role in the control of seed dormancy, and in mediating stress responses limiting the germination of non-dormant seeds (Kushiro *et al.*, 2004). We examined the expression of reporters associated with ABA response and synthesis over the time course of germination and compared this with the pattern in hydroprimed seeds to determine at which stage this treatment acted upon the ABA program.

The *RESPONSE TO ABA18* (*RAB18*) promoter has been characterized previously as responsive to ABA, and acts as a useful proxy to determine where ABA-mediated transcriptional responses are occurring (Lång and Palva, 1992). Over the early stages of germination, this reporter is principally concentrated in the radicle, with some activity present on the outer margins of the cotyledons (Figs. 4A-D). This pattern continues until the seed has completed germination (Figs 4E-H), where it then becomes focused first in the root of the early seedling (Fig. 4J) then the lower hypocotyl at a later stage of seedling development (Fig. 4K). Embryos from seeds which have been hydroprimed do not exhibit the radicle-enriched activity of the *RAB18::GUS* reporter (Fig 4L), but rather have this in the hypocotyl and cotyledons. The absence of a radicle-based pattern suggests the ABA response program has progressed beyond the early stages of the germination sequence. A similar pattern of radicle-enrichement is observed in the abundance of the ABA synthesis protein *ABA DEFICIENT 2* (*ABA2*) during the early stages of germination, and is absent in hydroprimed seeds (Supplementary Figure 6). The distribution of the ABA synthesis protein *ALDEHYDE OXIDASE 3* (*AAO3*) remains largely unchanged across the germination process and in hydroprimed embryos (Supplementary Figure 7).

**Figure 4.**
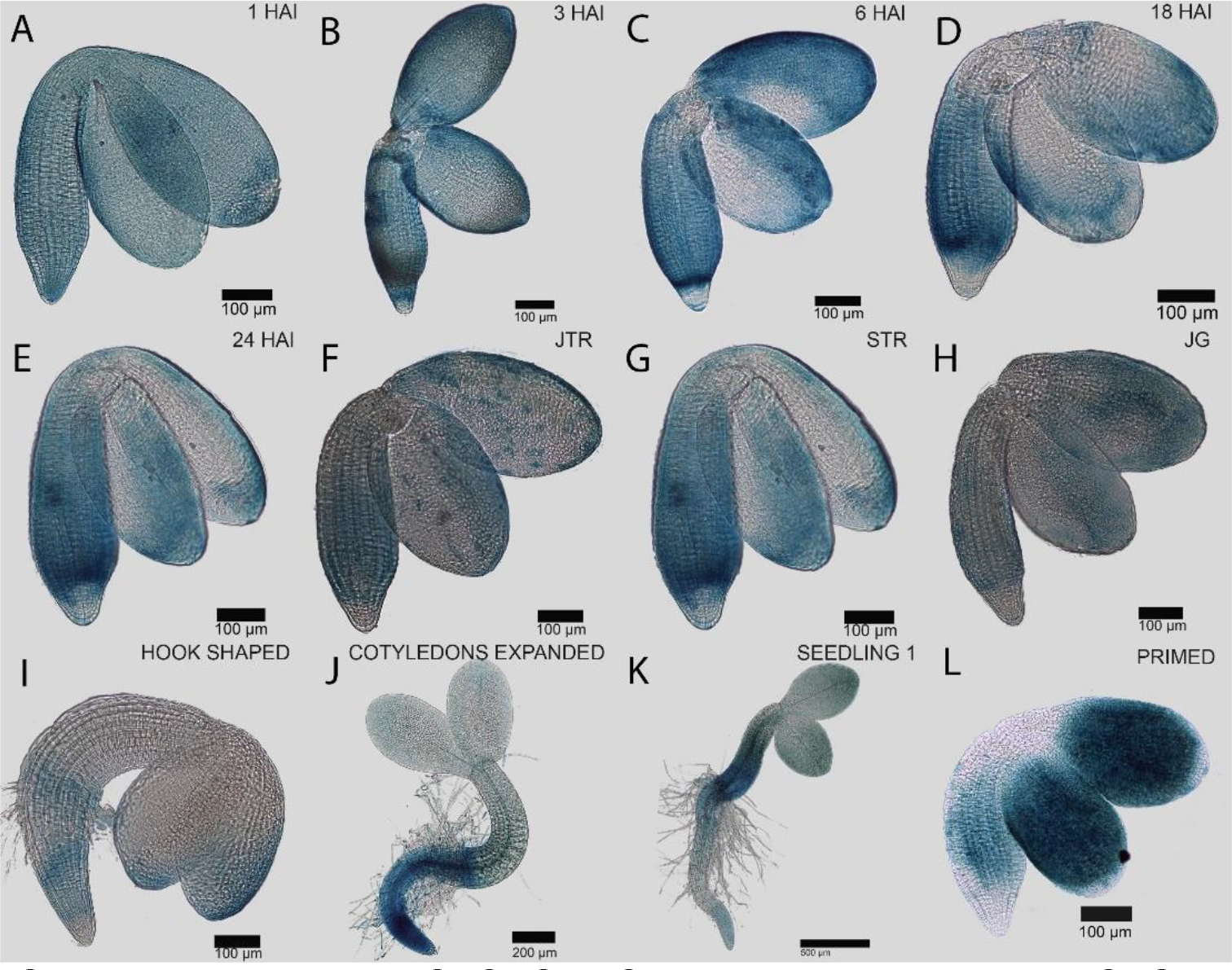
Spatial activity of the *RESPONSE TO ABA 18* (*RAB18*), *RAB18::GUS* promoter reporter in germinating *Arabidopsis* seeds and seedling establishment. GUS activity at an embryo at (A) 1 HAI, (B) 3 HAI, (C) 6 HAI, (D) 18 HAI, (E) 24 HAI, (F) early testa rupture, (G) late testa rupture, (H) immediately after germination, (I) hook shaped seedling stage, (J) expanded cotyledons in seedling and (K) fully established seedling. (L) Embryo from a seed that has been hydroprimed for 30 h at 22 oC. Black bars indicate the scale in each image.

These data suggest that ABA response and synthesis are not arrested at an early stage of the germination chronology, as is the case with gene expression associated with cell expansion.

### GA synthesis and signalling during the seed to seedling transition

GA synthesis is required for the induction of intact *Arabidopsis* seeds (Koornneef and Vanderveen, 1980). The patterns of gene expression associated with this induction of GA synthesis show cell type specific profiles (Ogawa *et al.*, 2003; Yamaguchi *et al.*, 2001), which are modulated by cold temperatures (Yamauchi *et al.*, 2004) and light (Yamaguchi *et al.*, 2001). This in turn leads to GA responses which act to promote downstream cell wall-associated gene expression in seeds (Cao *et al.*, 2006). Central to this induction are DELLA proteins (Lee *et al.*, 2002) and the *SCARECROW-LIKE3* (*SCL3*) transcription factor which controls germination responses (Zhang *et al.*, 2011). GUS reporters for GA synthesis, signalling and response components were also examined to understand the spatiotemporal events underpinning this hormone response which stimulates the germination process.

The *SCL3::GUS* reporter acts as a useful proxy to understand the cellular sites where GA responses are occurring (Zhang *et al.*, 2011). Activity of this reporter is enriched in the radicle during the early stages of seed germination (Figs 5A-E) and progressively moves along the hypocotyl and into the cotyledons until the completion of germination (Figs. 5F-H). Following germination, *SCL3::GUS* activity is enriched in the root (Figs 5I-K), and presumably in the endodermis where it has been reported previously to be present (Zhang *et al.*, 2011).

**Figure 5.**
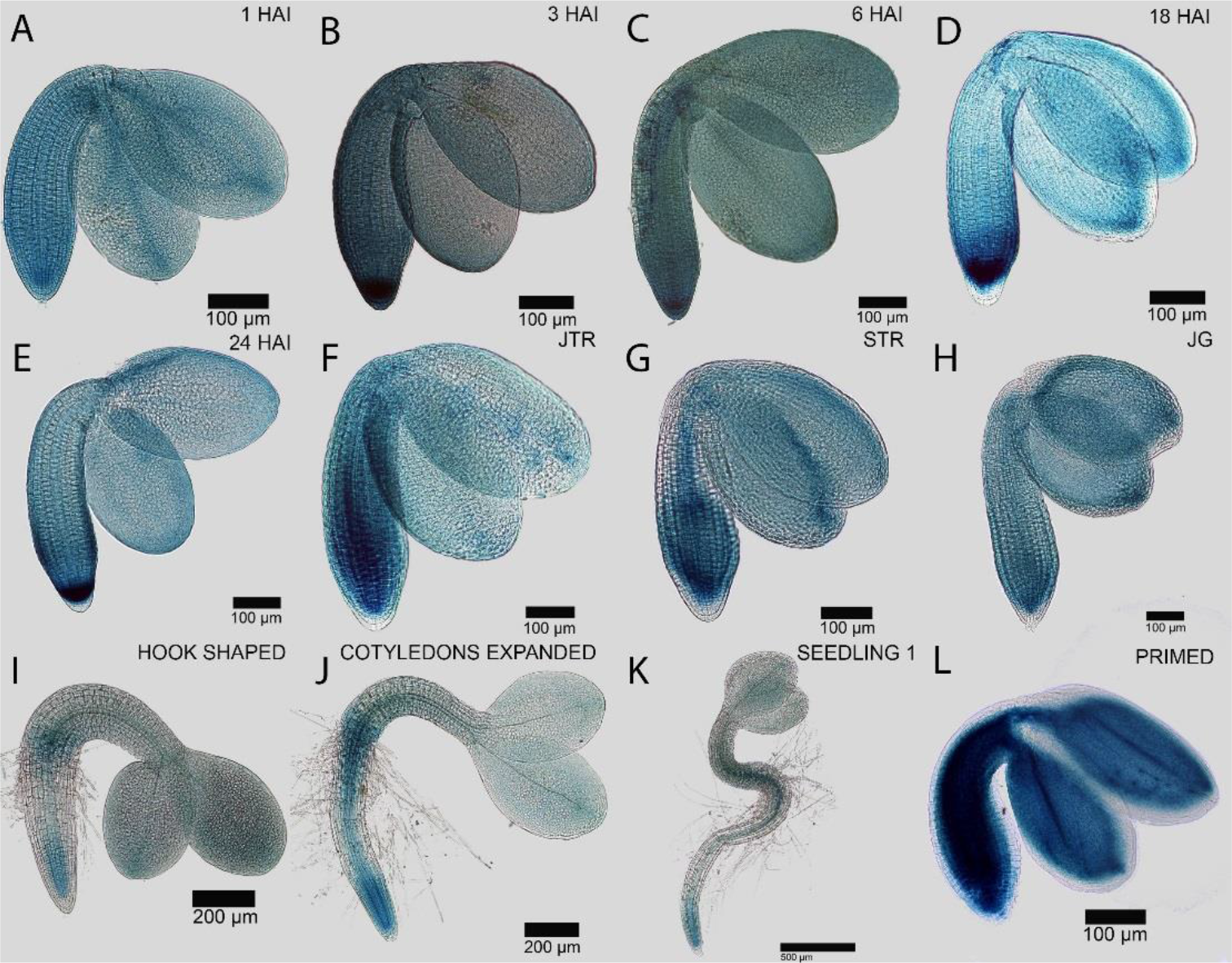
Spatial and temporal dynamics of the *SCL3::GUS* reporter during the seed to seedling transition in *Arabidopsis.* Promoter activity in the germinating embryo at (A) 1 HAI, (B) 1 HAI, (C) 3 HAI, (D) 6 HAI, (E) 18 HAI, (F) early testa rupture, (G) late testa rupture, and germinated seedlings just after the completion of germination (H), (I) hook stage seedling, (J) recently expanded cotyledons and (K) a fully established seedling. (L) Pattern of *SCL3::GUS* promoter activity in a hydroprimed embryo. Black bars indicate the scale in each image.

Embryos from hydroprimed seeds show a broad pattern of *SCL3::GUS* activity throughout the embryo similar to the later stages of germination (Fig. 5L). Notably, the activity of this reporter is absent from the epidermal cells of the embryo. This is in contrast to all other stages of the germination process, and suggests that the hydropriming process is differentially regulating the ability of the embryo to respond to GA specifically in the epidermis.

A similar phenomenon is present in the *GIBBERELLIN 3-OXIDASE 1* (*GA 3-OX1*) GUS reporter, representing the final and rate-limiting step of bioactive GA (Olszewski *et al.*, 2002). The initial induction of the *GA 3-OX1* promoter occurs broadly across the embryo radicle, and moves into the hypocotyl as germination progresses (Supplementary Figure 8). Hydroprimed seeds have broad activity of this promoter across the embryo axis, but is excluded from the epidermis (Supplementary Figure 8L). Conversely, the *GA 3OX2::GUS* reporter shows ectopic expression in embryos from hydroprimed seeds (Supplementary Figure 9L). The domain of GA synthesis is therefore not being globally restricted by hydropriming treatment.

The hormone GA is perceived by the *GIBBERELLIN INSENSITIVE DWARF* (*GID*) receptors (Ueguchi-Tanaka *et al.*, 2005). The presence of GID1A or GID1C proteins is a primary requirement for a cell to be able to respond to this hormone. We examined the distribution of these receptors using GUS translational fusions (Supplementary Figures 10-11), and found them to be broadly distributed across the germinating embryo. Hydropriming enriched GID1A to the vasculature (Supplementary Figure 10L) while GID1C remained broadly distributed, including in the embryo epidermis (Supplementary Figure 11L). The ability of the epidermis to perceive GA is therefore not limited by hydropriming treatment.

Collectively, these results indicate that hydropriming does not limit GA synthesis or perception in the embryo epidermis, but does limit the ability of these cells to actively respond to GA based on the absence of *SCL3::GUS* reporter activity.

### GA response in the embryo epidermis regulates hydropriming responses in Arabidopsis seeds

Given the differential expression of the GA-response marker *SCL3::GUS* in hydroprimed *Arabidopsis* embryos, we sought to investigate whether GA response in the epidermis impacts the response of seeds to this germination-enhancing treatment.

A previous study developed a transgenic *Arabidopsis* line specifically with altered GA responses in the epidermis (Rombolá-Caldentey *et al.*, 2014). By making use of the epidermis-specific At*ML1* promoter, the gain-of-function DELLA mutant protein *gai-1* was expressed in this cell type, including an N-terminal GFP tag (*ProML1:GFP-gai-1*).

These transgenic seeds with diminished GA responses specifically in the epidermis were hydroprimed along with non-transgenic control seeds (ecotype L*er*) to determine whether hormone response in this cell type impacted the germination-enhancing effects of this treatment.

Both the t10 and t95 showed little difference between the genotypes in hydroprimed seeds (Figs. 6A-B). The uniformity of germination was however dramatically different, with the *ProML1:GFP-gai-1* genotype showing less synchronous profiles across the population (Fig. 6C).

**Figure 6.**
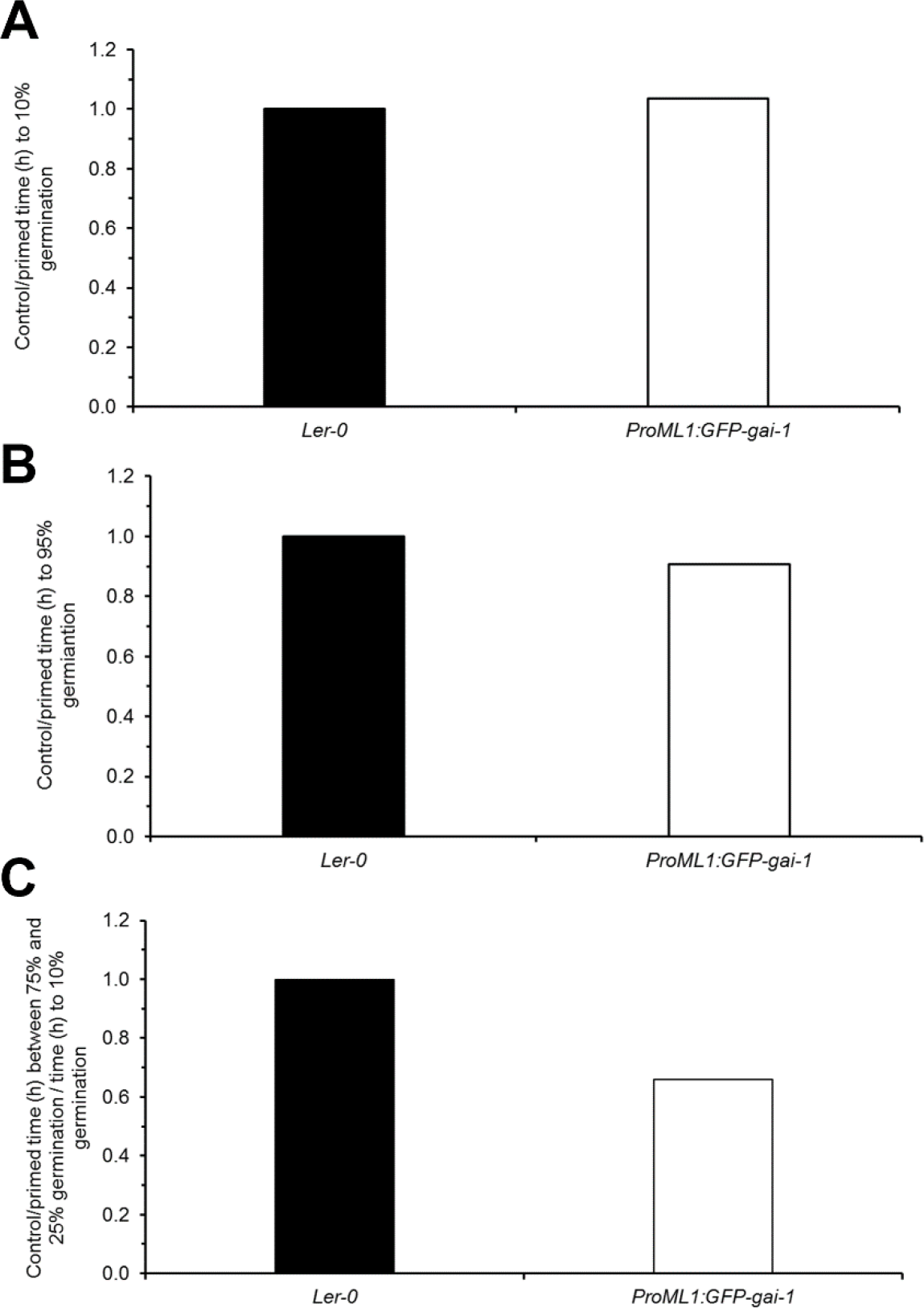
Impact of reduced GA response in the embryo epidermis on germination speed and uniformity following hydropriming. (A) Ratio of the time it took to reach 10% germination in control (L*er*) and transgenic *ProML1:GFP-gai-1* seeds before and after hydropriming. (B) Same as (A) for the time it took to reach 95% germination. (C) Same as (A) for the time it took to go from 25% to 75% germination. In all instances n = 3.

This result demonstrates that the control of GA response in the embryo epidermis of *Arabidopsis* is a key factor in controlling the ability of seeds to respond positively to hydropriming with response to germination uniformity.

## DISCUSSION

The sale of high quality seed lots having uniform germination underpins the $62.1 billion annual global seed trade. The application of treatments to seeds, including priming, are designed to increase the synchronization of the germination of these populations (Finch-Savage and Bassel, 2015). Due to the high cost of these treatments, they are typically limited to high value horticultural seeds and a limited number of field crops (Paparella *et al.*, 2015). Despite the economic and agronomic importance of seed priming, little is known as to how they operate at a mechanistic level. This study sought to understand where the seed priming treatment of hydropriming acts to limit germination, and in turn enhance the synchronization of seed lots. By characterizing the sequential steps underlying seed germination for gene expression associated with cell growth, and hormone synthesis and response, a molecular developmental chronology was established.

Coupling these data with hydropriming treatments for *Arabidopsis* seeds enabled the developmental point at which this process acts to limit germination. In the case of growth-promoting gene expression, which represents the downstream processes driving the germination process itself, these were blocked at the early stages of the program. By contrast, ABA and GA-associated molecular events progressed to later stages. This provides insight into the temporal regulation between the hormonal control of germination and the downstream processes which directly support this process.

In hydroprimed seeds, GA response was specifically impacted in the epidermis based on the absence of *SCL3::GUS* promoter activity in treated seeds (Fig. 5L). This combination of microscopy and molecular genetics revealed differential cell type specific control of underlying the priming response in *Arabidopsis* seeds. A central role for GA responses in promoting germination uniformity in response to hydropriming was provided using transgenic seed where GA responses are specifically diminished in the epidermis. This led to decreases in the uniformity of seed germination following hydropriming (Fig. 6C).

This has identified both a genetic pathway and cell type which act to promote germination uniformity in response to hydropriming. This provides both molecular and cellular targets which can be manipulated to enhance seed performance.

Heterogeneity in the germination of populations of seeds can arise from two major sources: the time at which the developmental fate switch to commence germination is flipped, and the rate at which the germination program proceeds (Finch-Savage and Bassel, 2015). In order to have uniformity in the germination of a population of seeds, the end point must be reached at the same time. In light of these two variables acting simultaneously, co-ordinating the timing of this transition endpoint proves problematic.

Based on our analyses, it appears that hydropriming acts to block germination at an early stage of this process. We therefore conclude that the mechanism of synchronization in this instance is through ensuring that all seeds have commenced the germination program, and are arrested at an early stage. The remaining heterogeneity in the germination of populations of hydroprimed seeds result from the rate at which the developmental program leading to the establishment of a seedling is executed.

This work sheds insight into the molecular and cellular basis of seed priming, providing targets for genetic manipulation to enhance seed quality independently of costly treatments.

## MATERIALS AND METHODS

### Plant growth conditions

All plants were grown in environmentally controlled cabinets, using 16 h light (23°C) and 8 h dark at 22 °C. Seeds were collected when plants had stopped flowering, and placed in glassine bags for 4 weeks to remove primary dormancy. Seeds were then cleaned by passing dried plant material through a fine mesh, and seeds were collected for use in subsequent experiments.

### Germination assays

Germination of *A. thaliana* seeds was conducted by plating 30 seeds in reps of 3. Seeds were scored for radicle emergence (germination) every 4 h until 100% germination was reached. All seeds were on sterilised with 10% bleach and germinated on ½ MS 0.8% (w/v) agar plates in a growth room conditions with a 16 hour light 8 hour dark photoperiod at 22°C.

### Hydropriming protocol

*A. thaliana* seeds were hydroprimed by submerging seeds in aerated sterile distilled water in nylon mesh bags for selected periods of time at either 10, 16 or 22 °C. Following priming drying was conducted by placing seeds between two layers or filter paper for 16 h before plating on ½ MS 0.8% (w/v) agar plates.

### Statistical analysis of germination uniformity

Statistical analysis of germination curves was conducted using the Germinator software package (Joosen *et al.*, 2010), to extract selected parameters; time taken to reach 10% germination (t10) 50% germination (t50), 95% germination (t95) and time between 25% and 75 (U7525) germination. The t10 parameter is considered as an indicator of dormancy, t50 represents the midpoint of the germination curve, t95 gives provides an indicator for total germination and U7525 is used as a uniformity measure.

### Reporter constructs generation

Reporter constructs for *EXPANSIN* genes were generated using 2 kb of sequence upstream of the ATG start codon for each gene as previously described (Bassel *et al.*, 2014). Other reporter constructs come from previous publications, including *XTH18* and *XTH19* (Vissenberg *et al.*, 2005b), *GID1A::GID1A-GUS* and *GID1C::GID1C-GUS* translational fusions (Suzuki *et al.*, 2009), *GA3ox1::GUS* and *GA3ox2::GUS* reporters (Hu *et al.*, 2008), *SCL3::GUS* (Zhang *et al.*, 2011), *AAO3::AAO3-GUS* and *ABA2::ABA2-GUS* (Seo *et al.*, 2006), and *RAB18::GUS* (Ghassemian *et al.*, 2000).

### GUS histochemical staining

*A. thaliana* embryos were dissected from seeds using a scalpel and forceps using a Leica SD6 microscope binocular microscope. Embryos were stained in X-Gluc solution with 0.1 M sodium phosphate buffer (pH 7.0), 0.1% Triton X-100 and 2 mM X-Gluc (Sigma). Embryos were stained at 37 °C until the blue substrate became visible, or for 24 h. Samples were fixed in a 3:1 ethanol/acetic acid, 500:1 DMSO, 1% Tween 20 fixative solution for 24 h and cleared in a chloral hydrate clearing solution until embryos were clear for imaging. Embryos were imaged using a Lecia DM500 light microscope

## ACKNOWLEDGEMENTS

I.S. was funded by summer studentship MIBTP BB/M01116X/1, N. K. M. was funded by a scholarship of Academic Training Scheme (SLAI) from the Ministry of Education Malaysia, Faculty of Agro Based Industry, University Malaysia Kelantan, Jeli Campus, and G.W.B. was funded by BBSRC grants BB/L010232/1, BB/J017604/1, and BB/N009754/1, Leverhulme grant RPG-2016-049.

## SUPPLEMENTARY FIGURE LEGENDS

**Supplementary Figure 1.** Establishment of a hydropriming protocol for *Arabidopsis* seeds. Time course of germination percentage for seeds hydroprimed at (A) 10 oC, (B) 16 °C, and (C) 22 °C for 6 h, 18 h, 30 h, and 42 h. (D) Time it took seeds to each 10% germination (t10) for each of the treatments in (A)–(C) expressed as a ratio of h in control seed over h in primed seeds . (E) Time it took seeds to each 50% germination (t50) for each of the treatments in (A)–(C) expressed as a ratio of h in control seed over h in primed seeds. (E) Time it took seeds to each 95% germination (t95) for each of the treatments in (A)–(C) expressed as a ratio of h in control seed over h in primed seeds. (F) Uniformity in germination for each of the treatments based on time taken between achieving 25% and 75% germination. Error bars in (A)-(C) are SD with n = 3.

**Supplementary Figure 2.** Spatial activity of the *EXPANSIN 9*, *EXPA9::GUS* promoter reporter in germinating *Arabidopsis* seeds and seedling establishment. GUS activity at an embryo at (A) 1 HAI, (B) 3 HAI, (C) 6 HAI, (D) 18 HAI, (E) 24 HAI, (F) early testa rupture, (G) late testa rupture, (H) immediately after germination, (I) hook shaped seedling stage, (J) expanded cotyledons in seedling and (K) fully established seedling. (L) Embryo from a seed that has been hydroprimed for 30 h at 22 °C. Black bars indicate the scale in each image.

**Supplementary Figure 3.** Spatial activity of the *EXPANSIN 10*, *EXPA10::GUS* promoter reporter in germinating *Arabidopsis* seeds and seedling establishment. GUS activity at an embryo at (A) 1 HAI, (B) 3 HAI, (C) 6 HAI, (D) 18 HAI, (E) 24 HAI, (F) early testa rupture, (G) late testa rupture, (H) immediately after germination, (I) hook shaped seedling stage, (J) expanded cotyledons in seedling and (K) fully established seedling. (L) Embryo from a seed that has been hydroprimed for 30 h at 22 °C. Black bars indicate the scale in each image.

**Supplementary Figure 4.** Spatial activity of the *EXPANSIN 15*, *EXPA15::GUS* promoter reporter in germinating *Arabidopsis* seeds and seedling establishment. GUS activity at an embryo at (A) 1 HAI, (B) 3 HAI, (C) 6 HAI, (D) 18 HAI, (E) 24 HAI, (F) early testa rupture, (G) late testa rupture, (H) immediately after germination, (I) hook shaped seedling stage, (J) expanded cotyledons in seedling and (K) fully established seedling. (L) Embryo from a seed that has been hydroprimed for 30 h at 22 °C. Black bars indicate the scale in each image.

**Supplementary Figure 5.** Spatial activity of the *XYLOGLUCAN ENDOTRANSGLUCOSYLASE/HYDROLASE 18*, *XTH18::GUS* promoter reporter in germinating *Arabidopsis* seeds and seedling establishment. GUS activity at an embryo at (A) 1 HAI, (B) 3 HAI, (C) 6 HAI, (D) 18 HAI, (E) 24 HAI, (F) early testa rupture, (G) late testa rupture, (H) immediately after germination, (I) hook shaped seedling stage, (J) expanded cotyledons in seedling and (K) fully established seedling. (L) Embryo from a seed that has been hydroprimed for 30 h at 22 °C. Black bars indicate the scale in each image.

**Supplementary Figure 6.** Spatial activity of the *ABA DEFICIENT 2* (*ABA2*), *ABA2::ABA2-GUS* translational fusion reporter in germinating *Arabidopsis* seeds and seedling establishment. GUS activity at an embryo at (A) 1 HAI, (B) 3 HAI, (C) 6 HAI, (D) 18 HAI, (E) 24 HAI, (F) early testa rupture, (G) late testa rupture, (H) immediately after germination, (I) hook shaped seedling stage, (J) expanded cotyledons in seedling and (K) fully established seedling. (L) Embryo from a seed that has been hydroprimed for 30 h at 22 °C. Black bars indicate the scale in each image.

**Supplementary Figure 7.** Spatial activity of the *ALDEHYDE OXIDASE 3* (*AAO3*), *AAO3::AAO3-GUS* translational fusion reporter in germinating *Arabidopsis* seeds and seedling establishment. GUS activity at an embryo at (A) 1 HAI, (B) 3 HAI, (C) 6 HAI, (D) 18 HAI, (E) 24 HAI, (F) early testa rupture, (G) late testa rupture, (H) immediately after germination, (I) hook shaped seedling stage, (J) expanded cotyledons in seedling and (K) fully established seedling. (L) Embryo from a seed that has been hydroprimed for 30 h at 22 °C. Black bars indicate the scale in each image.

**Supplementary Figure 8.** Spatial activity of the *GIBBERELLIN 3 G-HYDROXYLASE 1* (*GA3ox1*), *GA3ox1::GUS* promoter reporter in germinating *Arabidopsis* seeds and seedling establishment. GUS activity at an embryo at (A) 1 HAI, (B) 3 HAI, (C) 6 HAI, (D) 18 HAI, (E) 24 HAI, (F) early testa rupture, (G) late testa rupture, (H) immediately after germination, (I) hook shaped seedling stage, (J) expanded cotyledons in seedling and (K) fully established seedling. (L) Embryo from a seed that has been hydroprimed for 30 h at 22 °C. Black bars indicate the scale in each image.

**Supplementary Figure 9.** Spatial activity of the *GIBBERELLIN 3 G-HYDROXYLASE 2* (*GA3ox2*), *GA3ox2::GUS* promoter reporter in germinating *Arabidopsis* seeds and seedling establishment. GUS activity at an embryo at (A) 1 HAI, (B) 3 HAI, (C) 6 HAI, (D) 18 HAI, (E) 24 HAI, (F) early testa rupture, (G) late testa rupture, (H) immediately after germination, (I) hook shaped seedling stage, (J) expanded cotyledons in seedling and (K) fully established seedling. (L) Embryo from a seed that has been hydroprimed for 30 h at 22 °C. Black bars indicate the scale in each image.

**Supplementary Figure 10.** Spatial activity of the *GIBBERELLIN INSENSTIVIE DWARF 1A* (*GID1*), *GID1A::GID1A-GUS* translational fusion reporter in germinating *Arabidopsis* seeds and seedling establishment. GUS activity at an embryo at (A) 1 HAI, (B) 3 HAI, (C) 6 HAI, (D) 18 HAI, (E) 24 HAI, (F) early testa rupture, (G) late testa rupture, (H) immediately after germination, (I) hook shaped seedling stage, (J) expanded cotyledons in seedling and (K) fully established seedling. (L) Embryo from a seed that has been hydroprimed for 30 h at 22 °C. Black bars indicate the scale in each image.

**Supplementary Figure 11.** Spatial activity of the *GIBBERELLIN INSENSTIVIE DWARF 1C* (*GID1C*), *GID1C::GID1C-GUS* translational fusion reporter in germinating *Arabidopsis* seeds and seedling establishment. GUS activity at an embryo at (A) 1 HAI, (B) 3 HAI, (C) 6 HAI, (D) 18 HAI, (E) 24 HAI, (F) early testa rupture, (G) late testa rupture, (H) immediately after germination, (I) hook shaped seedling stage, (J) expanded cotyledons in seedling and (K) fully established seedling. (L) Embryo from a seed that has been hydroprimed for 30 h at 22 °C. Black bars indicate the scale in each image.

